# The build-up of the present-day tropical diversity of tetrapods

**DOI:** 10.1101/2022.12.05.519156

**Authors:** Ignacio Quintero, Michael Landis, Walter Jetz, Hélène Morlon

## Abstract

The extraordinary number of species in the tropics when compared to the extra-tropics is probably the most prominent and consistent pattern in biogeography, suggesting that overarching processes regulate this diversity gradient. A major challenge to characterizing which processes are at play relies on quantifying how the frequency and determinants of tropical and extra-tropical speciation, extinction and dispersal events shaped evolutionary radiations. We address this question by developing and applying spatio-temporal phylogenetic and paleontological models of diversification for tetrapod species incorporating paleoenvironmental variation. Our phylogenetic model results show that area, energy or species richness did not uniformly affect speciation rates across tetrapods and dispute expectations of a latitudinal gradient in speciation rates. Instead, both neontological and fossil evidence coincide in underscoring the role of extra-tropical extinctions and the outflow of tropical species in shaping biodiversity. These diversification dynamics accurately predict present-day levels of species richness across latitudes and uncover temporal idiosyncrasies but spatial generality across the major tetrapod radiations.

## Main Text

The present-day increase in species diversity towards the tropics is one of the most widespread biogeographical patterns, shared by a wide array of taxa, including microorganisms, fungi, insects, plants, and vertebrates (Willig et al., 2003). The generality of this latitudinal diversity gradient (LDG) across continents and clades suggests that universal spatial and temporal evolutionary processes transcend the idiosyncratic ecologies of different lineages (Rohde, 1992; Mittelbach et al., 2007). Since differences in species richness result from differences in speciation, dispersal and extinction events, any mechanisms explaining the LDG must ultimately link to variation on these processes (Wiens and Donoghue, 2004). Therefore, identifying general explanations for the LDG demands, first, to assess whether the relative contributions among these evolutionary processes coincide across taxa and, second, to investigate if the mechanisms behind this variation are shared.

How speciation, dispersal, and extinction cause tropical and extra-tropical species richness to differ has been widely debated. On one hand, variation in species richness can result from differences in evolutionary rates (Mittelbach et al., 2007; Morlon, 2020). For instance, the tropics could be characterized by abiotic and biotic conditions favorable to speciation, persistence (including lower extinction rates) and/or higher immigration rates when compared to the extratropics (Wiens and Donoghue, 2004; Jablonski et al., 2006). On the other hand, an older origin and persistence in the tropical biome could provide more time for species to accumulate, even in the absence of rate differences (Mittelbach et al., 2007). Studies integrating information from both the recent and deeper past generally supported the ‘evolutionary rate’ hypothesis, with evidence for higher speciation rates in species-rich areas (Mittelbach et al., 2007). In contrast, several newer studies using recent speciation rates found either no association with richness (Jetz et al., 2012; Belmaker and Jetz, 2015) or markedly higher rates in depauperate areas (associated with higher elevations/latitudes) (Quintero and Jetz, 2018; Rabosky et al., 2018; Igea and Tanentzap, 2020; Harvey et al., 2020), advancing the possibility that tropical relative to extratropical speciation rates have changed from past to present (Schluter and Pennell, 2017; Morlon, 2020). Rigorously testing these hypotheses, as well as identifying which factors influence temporal variation in tropical relative to extra-tropical speciation rates, have been hampered by the lack of diversification models that account for spatio-temporal evolutionary dynamics in a changing environment and that test for congruence between neontological and paleontological evidence (Fritz et al., 2013). Time-calibrated phylogenetic trees built from the genetic data of extant contain topological and temporal information that, through models, indirectly informs how speciation operates. Fossil information, on the other hand, remains the only direct evidence of extinction and past species distributions, yet its incompleteness and fragmentary nature hinders a comprehensive characterization of species relationships and their branching process for most clades (Behrensmeyer et al., 2000; Silvestro et al., 2016). Approaches using simulation tools have been useful (Rangel et al., 2018; Hagen et al., 2021), but these frameworks do not provide inference methods to estimate evolutionary parameters from empirical datasets.

We first developed a new set of phylogenetic models that account for the effect of paleoenvironmental variations and observed as well as unobserved (‘hidden’) species states on spatial diversification, which we call ‘ESSE’ (Materials and Methods & Supplementary Information). These models allow diversification rates to vary in space and time while accounting for environmental fluctuations and variation that is independent of the geographical distribution of lineages – for example, in relation to other ecological attributes of the species. Our hypothesis testing controls for background rate variation that is independent of the tropical and/or extra-tropical identity of lineages through a hidden-states framework (Beaulieu and O’Meara, 2016). We explored the latitudinal diversification dynamics behind the build-up of the present-day LDG by combining this model with spatial and phylogenetic information for 32,791 tetrapod taxa, representing most mammal, bird, amphibian and squamate extant species (Materials and Methods). We delineate the tropical biome from the extra-tropics following a biologically relevant environmental definition based on the Köppen climate classification (Beck et al., 2018)(Fig 1a). This definition captures the high productivity and energy characterizing the tropical biome and avoids complications when using other overly inclusive definitions, such as the 23.5° latitudinal cut-off that lumps contrasting biomes like deserts together with evergreen forests. Collating range distribution information at the validated *ca*. 110km resolution for all species (Hurlbert and Jetz, 2007), we assigned each species as either tropical, extra-tropical or widespread. Consistent with previous studies, our data display a strong latitudinal gradient in species richness across the four major tetrapod clades (Fig 1b) and concomitant richness differences between the tropics and the extra-tropics (Fig 1d). Our analyses used time-calibrated phylogenetic trees with complete species-level sampling for each of the tetrapod clades; species that were non-genetically represented were pruned out but accounted for using regionally-specific sampling fractions (Materials and Methods) to avoid potential inference biases caused by the uncertain placement of these lineages (Rabosky, 2015).

**Figure 1:**
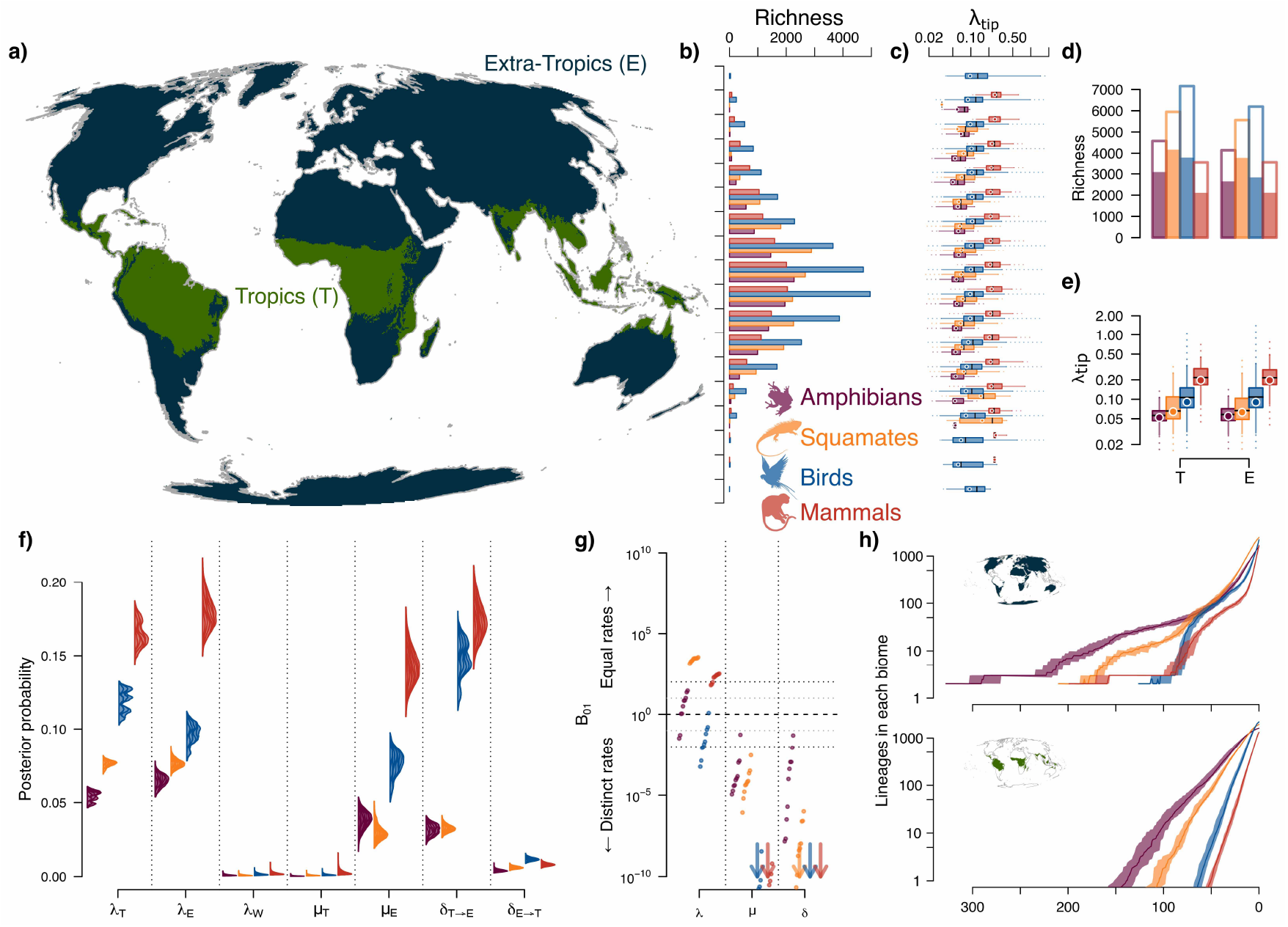
Latitudinal tetrapod richness and diversification patterns. **a)** Map of tropical (T) and extra-tropical (E) delimitation. **b-e)** Total tetrapod richness and their present-day speciation rates (*λ*_tip_, an average over 10 trees) distributions aggregated by **b-c)** 10° wide latitudinal bands and by **d-e)** tropical and extra-tropical regional delimitation (endemics are shown here as solid bars). Circles represent the median weighted by species range size. **f)** Stacked posterior distributions (with density standardized across all parameters for visualization) across 10 trees for each tetrapod clade for speciation in the tropics (*λ*_*T*_), extra-tropics (*λ*_*E*_), and widespread lineages (*λ*_*W*_), extinction in the two biomes (*μ*_*T*_ *& μ*_*E*_) and dispersal (*δ*) from the tropics to the extra tropics (T → E) and vice versa (E → T) for model ‘G’. **g)** Bayes factors (BF) between constant or different rates in speciation, extinction and dispersal (an arrow shows the direction if one or more BFs exceed *>* 10^10^ in magnitude). **h)** Lineage Through Time (LTT) plot for each biome as the sum of the posterior marginal ancestor state probabilities through time.

### Elevated extratropical extinction maintains diversity gradient

We explored geographic patterns of diversification, first ignoring potential temporal and hidden heterogeneity (model ‘G’; Materials and Methods). In light of inferential concerns about diversification models (Louca and Pennell, 2020, 2021), we stress that our models are identifiable, and we show that biogeographic information strengthens the confidence we can have in extinction rate estimates (see Supplementary Information “Macroevolutionary Rates”). Contrary to previous studies integrating deep timescales on previous biological datasets that suggest a negative latitudinal trend with origination rates (Jablonski et al., 2006; Mittelbach et al., 2007; Rolland et al., 2014) and to studies focusing on recent speciation rates that suggest a positive latitudinal trend (Weir and Schluter, 2007; Rabosky et al., 2018; Igea and Tanentzap, 2020; Harvey et al., 2020), and matching findings for birds (Jetz et al., 2012), we find that speciation rates have been remarkably similar between the tropics and the extra-tropics across the four major tetrapod radiations, regardless of whether we focus on recent speciation rates (Fig 1c,e) or integrate deeper timescales (Fig 1f,g). Bayesian model selection supported models with equal speciation rates between tropics and extra-tropics for most clades (except slightly higher tropical rates in birds, Fig 1f,g). Post-hoc association tests of tip speciation rates with either latitudinal bands (Fig 1c) or aggregated tropical or extra-tropical taxa (Fig 1e) were far from significant (Fig S1). Our simplest model therefore rejects the hypothesis of higher speciation rates in the tropics relative to the extra-tropics. Similarly, our results do not support the idea that the abundant tropical richness results from more available time for speciation (Mittelbach et al., 2007): using ancestral states estimation, we find an extra-tropical origin among lineages that survived to the present across all clades followed by more recent colonization of the tropics (Fig 1h), in agreement with recovered crown fossils for amphibians (Fröbisch et al., 2010), squamates (Simoes et al., 2018), birds (Field et al., 2020) and mammals (Benson et al., 2013). Rather, our results suggest that substantially lower extinction rates gave rise to the extraordinary tetrapod richness of the tropics (Fig 1f), with all clades strongly supporting models with biome-dependent extinction rates (Fig 1g). In turn, extra-tropical richness suffers from higher extinction rates but is bolstered by a highly supported asymmetrical influx of tropical species (Fig 1f,g). These elevated rates of extra-tropical colonization by tropical lineages conflict with the ‘tropical niche conservatism’ idea (Wiens and Donoghue, 2004) but partly agree with the ‘Out of the Tropics’ hypothesis, wherein tropical clades steadily colonize the extra-tropics over time (Jablonski et al., 2006).

Incorporating hidden states to these geographical models (model ‘G+H’), reveal that tropical and extra-tropical lineages each separate into two further diversification regimes (Caetano et al., 2018) (Fig 2a,b; Materials and Methods). Regardless of the hidden state (0 or 1; *i*.*e*., biome-independent variation), the tropics act as a ‘source’, accumulating species steadily with a net outflow of species towards the extra-tropics. In contrast, the extra-tropics either experience more extinction than speciation events, acting as a ‘sink’, or experience more speciation than extinction events, depending on the lineages’ hidden state (Fig 2a,b). Among the high influx of lineages towards the extra-tropics, our model detects which lineages became evolutionarily successful and which were more prone to rapid extinction. The lineages with high recent extratropical speciation rates correspond to those with a deep-time net diversification rate comparable to that in the tropics (Fig 2c,d; strongly supported associations, Table S2), challenging the idea that these fast speciating lineages are ephemeral on account of their high extinction rates (Quintero and Jetz, 2018; Harvey et al., 2020). These lineages also have a present-day net diversification rate that does not exceed tropical rates (Figs 1e,2a), undermining the possibility that a rapidly diversifying present-day extra-tropical biota could eventually achieve comparable diversity to the tropics (Rabosky et al., 2018).

**Figure 2:**
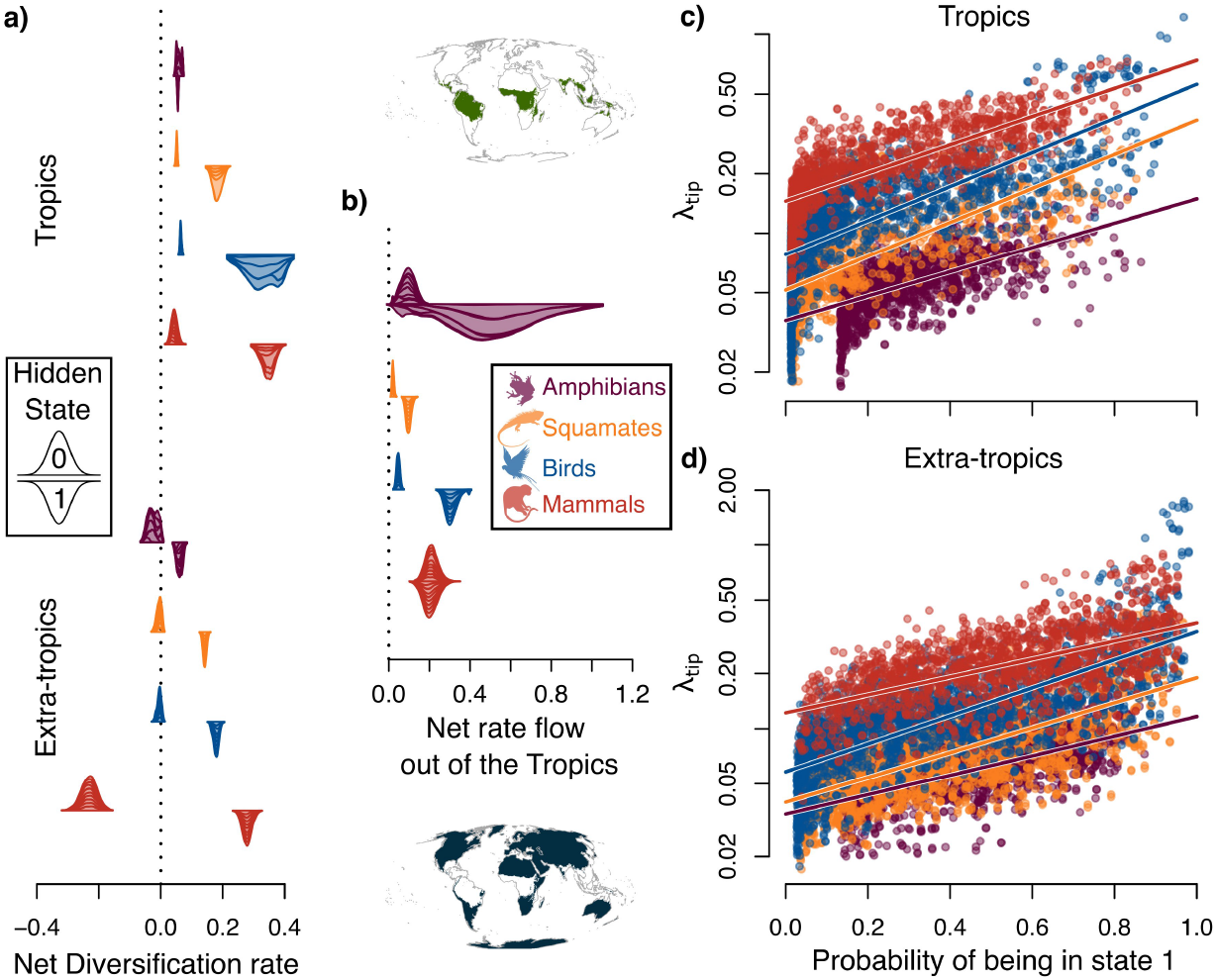
Heterogeneity in diversification rates within biomes. **a-b)** Posterior distributions (with density standardized across all parameters for visualization) across 10 trees for each tetrapod clade for **a)** net diversification rates (*λ* − *μ*) in the tropics (top) and extra-tropics (bottom) and for **b)** net rate flow out of the Tropics (*δ*_(T→E)_ − *δ*_(E→T)_), for two hidden states (0: up-facing and 1: down-facing, representing rate unobserved differences that are independent of the biome) for model ‘G+H’. **c-d)** Marginal posterior tip probabilities of being in state 1 versus corresponding tip-speciation rate (*λ*_tip_) for **c)** tropical and **d)** extra-tropical species. Clade colours as in Fig 1.

### Environmental change elicits idiosyncratic diversification responses across clades

To test the hypothesis that the relation of tropical to extra-tropical speciation rates have changed from past to present and to identify factors that have potentially influenced this temporal variation, we consider a series of ecologically important variables that have changed through time and are thought to influence speciation rates. How total terrestrial tropical and extra-tropical area has fluctuated dramatically across geological history (Mannion et al., 2014)(Fig 4a,b) influences phylogenetic model inferences (Landis et al., 2022), where speciation rates are predicted to be higher in large geographical regions by offering more opportunities of within-area population divergence and by supporting species with wider ranges (Kisel and Barraclough, 2010; Jetz and Fine, 2012). Similarly, area expansion can increase speciation rates as lineages track their abiotic preferences across space, inducing geographical range shifts and fragmentation (Barnosky, 2001). Temperatures have also fluctuated across geological history (Fig 3c), and warmer environments are expected to enhance speciation by being more productive environments that support higher population numbers and by accelerating evolutionary speed at the molecular level, offering more opportunities for population differentiation (Rohde, 1992; Allen et al., 2002). Speciation rates might also increase with the rate of temperature change, through climate-driven dispersal and vicariance events that isolate populations (Barnosky, 2001). Lastly, latitudinal differences in resource availability could either constrain speciation rates under a negative diversity-dependence scenario (Rabosky, 2013) or promote speciation by increasing the probability of divergence either by interspecific competition or by lowering average population numbers, and by generating higher community complexity through a ‘diversity begets diversity’ scenario (Emerson and Kolm, 2005).

**Figure 3:**
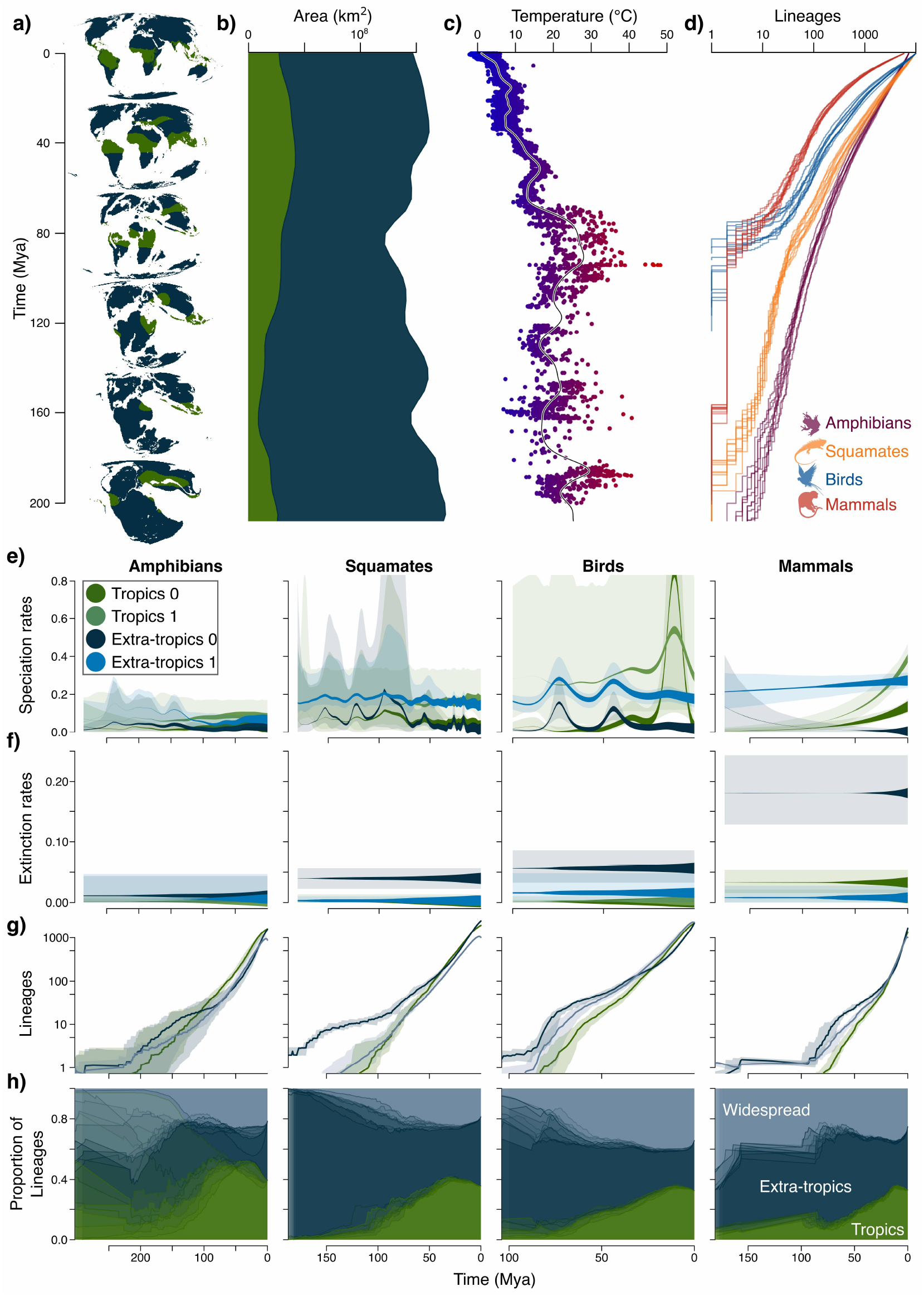
Spatio-temporal diversification dynamics in tetrapods. **a)** Maps every 40 My of tropical and extratropical terrestrial area. **b)** Tropical and extra-tropical area reconstruction, **c)** temperature and **d)** LTT plots for the last 200 My. **e)** Time-varying speciation and **f)** time-constant extinction rates and for each of the two hidden states for the tropics and the extra-tropics with 95% HPD uncertainty as background shade using model averaging by weighting each tree and parameters contributions by their posterior probability. The thickness of the line corresponds to the state-specific reconstructed number of lineages (in natural logarithm) that survived to the present (model for model ‘G+E+H’). **g)** Biome-specific LTT aggregated from ancestral posterior probabilities of being tropical, extra-tropical or widespread. **h)** LTT proportions in each state across time.

**Figure 4:**
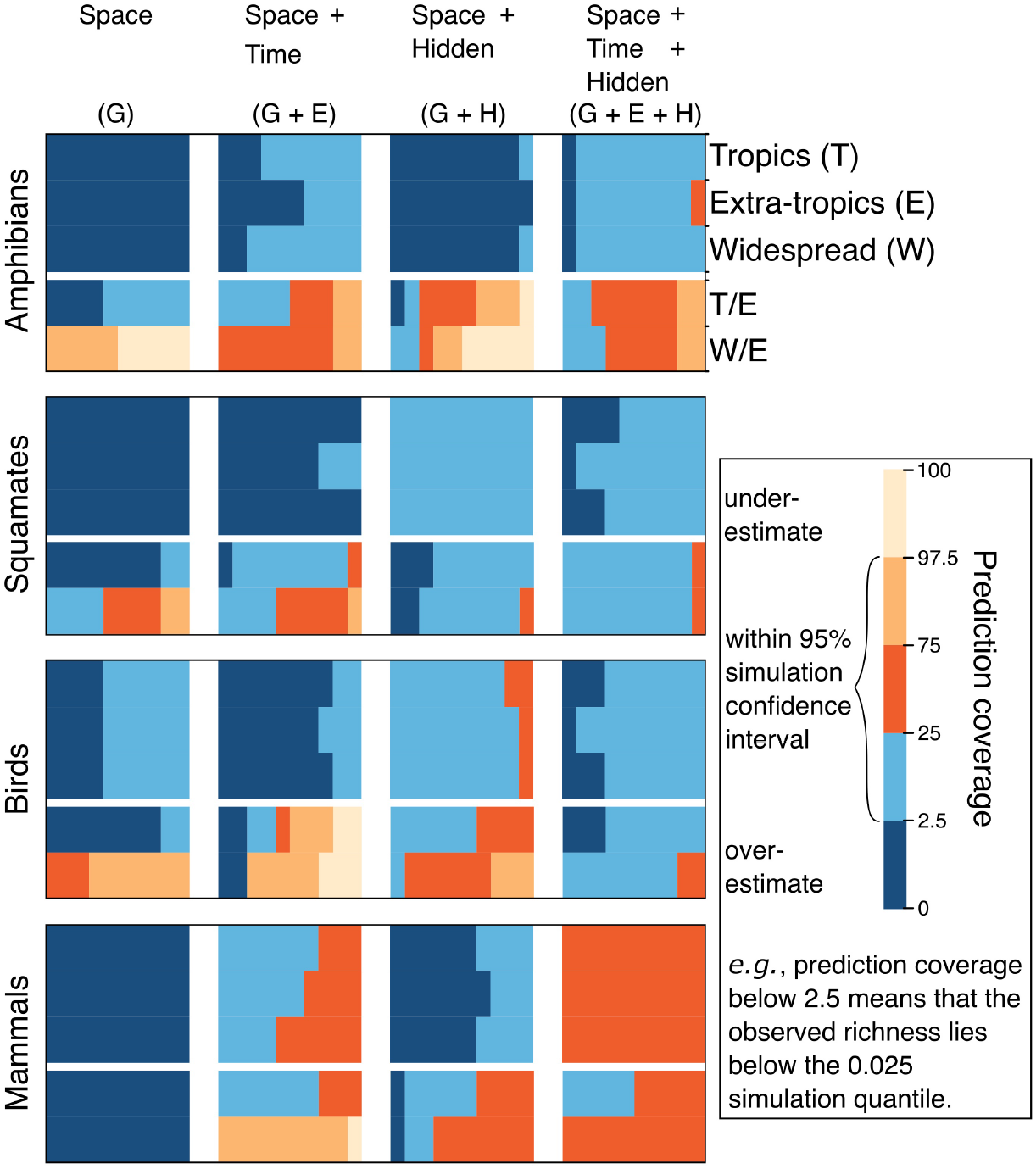
Goodness of fit of diversification models. Prediction coverage from the richness probability distributions for diversification models, in increasing complexity: Space = geographic dependence (G), Time = environmental dependence (E) and Hidden = hidden states (H). Prediction coverage is defined as the model-predicted quantile of species richness using simulations that contains the empirical measurement of true species richness. For each clade (rows), for each model (column), a subrow consists of results for each of the 10 trees in sequence (subcolumns), and represent, in order, tropical, extra-tropical and widespread richness and proportion of tropical/extratropical and widespread/extra-tropical.

We tested if regional species richness is explained by speciation rates controlled by habitat-specific area and its rate of change, temperature and its rate of change, or diversity-dependence using a time-varying spatio-temporal model of lineage speciation, extinction, and dispersal (model ‘G+E+H’; Materials and Methods). These models are identifiable, as we allow rate heterogeneity across lineages and constrain the functional form of the rate dependencies (Supplementary Information). This process-based approach allows to better evaluate ecological theories underlying richness variation by integrating deep-time environmental covariates and spatial diversification dynamics rather than by correlating present-day covariates with extant biodiversity. We estimated tropical and extra-tropical terrestrial area by merging reconstructed Köppen biomes and Digital Elevation Models to reconstruct world-wide tropical and extra-tropical regions every 5 My for the past 540 Mya (Scotese, 2021) (Materials and Methods, Video S1), and used these reconstructions as region specific covariates of tropical and extra-tropical speciation rates, respectively (Fig 4b). Similarly, we compiled global temperature data for the past 526 Mya (Fig 4c) but allowed for different associations between tropical and extra-tropical regions with temperature. We used a regionally-specific exponential relationship of speciation rate with time to approximate the effect of diversity-dependence (Rabosky and Lovette, 2008) (Materials and Methods). We assumed regionally-specific, but time-constant, extinction rates as our model validation confirmed previously-noted difficulties in recovering time-varying extinction dynamics from phylogenies of present-day species (Lewitus and Morlon, 2018; Louca and Pennell, 2020) (see Supplementary Information). We show, however, that under simulated scenarios using empirical extinction rate curves we are able to recover the pertinent time-averaged extinction rates between regions (Supplementary Information).

Intriguingly, while we find that the correspondence across tetrapods in geographic diversification and dispersal patterns holds after incorporating temporal dynamics, the temporal dynamics themselves are highly idiosyncratic (model ‘G+E+H’, Fig 4e, Figs S7, S8 & S9). While time-averaged rates show similar patterns as the constant rate models (*i*.*e*., to models ‘G’ & ‘G+H’; Figs S8 & 9), we find support for distinct nonlinear speciation rate dynamics where the tropical and extra-tropical rates exceeded each other at different times in the past (Fig 4e). For amphibians and squamates different time dependencies on speciation were selected for different posterior trees, indicating that, while speciation has not been constant, the data do not tell us which time-dependent hypothesis is the most likely (Table S3). Model-averaged rate curves, however, show that tropical and extra-tropical speciation rates both decline in the recent past. Bird speciation rate dynamics most closely responded to the rate of area-change through time, with lineages experiencing higher speciation rates when area expands rapidly in the tropics and when area contracts rapidly in the extra-tropics (Fig 4e, Fig S9). For mammals, diversity-dependent speciation rates were strongly supported, but with opposite effects for the two regions. In the tropics, mammal diversity begets diversity regardless of species hidden state; in the less productive extra-tropics, diversity either has no strong effect or impedes diversity, which in combinations with high extinction rates, induce the ‘sink’ regime (Fig 4e,f). To test our model’s ability to predict present-day tetrapod latitudinal diversity patterns, we performed forward-time simulations under our fitted models to compare predicted and observed tropical and extra-tropical species richness (Fig S10, Materials and Methods). We found that, except for birds, models without regionally-independent rate heterogeneity in the diversification process (without hidden states) or that ignore temporal speciation dynamics consistently overestimate the number of species at present across regions and overpredict the ratio of widespread to extra-tropical lineages (Fig 4). Instead, models with temporal and geographical dependencies, coupled with region-independent rate heterogeneity, accurately predict the observed richness in number of tropical, extra-tropical and widespread species for all clades (Fig 4).

### Elevated extratropical extinction reflected in fossil data

For a comprehensive assessment of these spatial diversification dynamics, we applied an analogous biogeographic model that uses fossil occurrences instead of phylogenetic trees, ‘DES’, and that accounts for several sources of uncertainty such as temporal and spatial sampling and preservation (Silvestro et al., 2016). While existing paleontological biogeographic models do not infer speciation rate dynamics (in ‘DES’ the processes of origination and range inheritance are ignored since it assumes no underlying phylogenetic structure across fossil data (Silvestro et al., 2016)), they provide direct observations of past extinction and dispersal events and relax the assumption of constant extinction rates through time for each area, evaluating the robustness of our phylogenetic models results. We focused on mammal dynamics during the Cenozoic since they have by far the best paleontological record of a terrestrial clade across tropical and extra-tropical regions (Mannion et al., 2014; Davies et al., 2017), and carefully vetted spatial fossil occurrence data at the genus level, following standard paleontological practice (Materials and Methods). We then allocated fossil occurrences using their timing and paleocoordinates to the spatial reconstructions of global tropical and extra-tropical regions.

Remarkably, our paleontological results concur with those from our phylogenetic methods, in that the extra-tropical region experienced higher extinction rates together with higher immigration of tropical diversity, regardless of assumptions of temporal rate constancy or heterogeneity across separate stratigraphic periods, while accounting for temporal and spatial fossil sampling differences (Fig 5). While the use of different taxonomic resolutions as well as different taxonomic concepts (Archibald, 1993; Silvestro et al., 2018) make phylogenetic and fossil based model rates not directly comparable, they highlight the relative region-specific rate differences in extinction and dispersal (Silvestro et al., 2016). Our results using phylogenetic information are also congruent with other results based on paleontological evidence (Mannion et al., 2014; Saupe et al., 2019), highlighting the utility of combining biogeographic and birth-death information for capturing past diversification dynamics.

**Figure 5:**
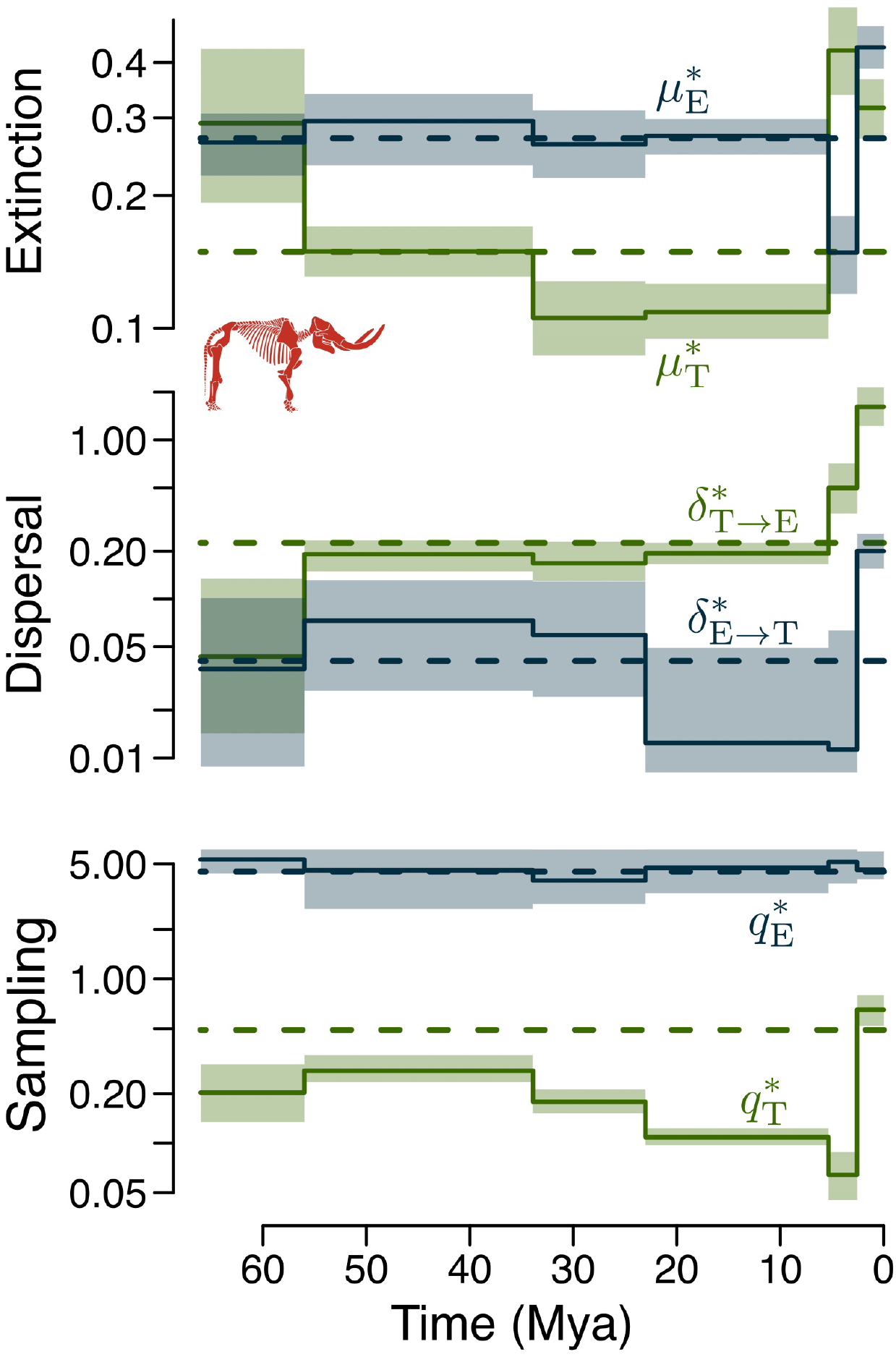
Spatio-temporal diversification dynamics in mammal fossil record. Results from Cenozoic fossil biogeographic analyses at the genus level for tropics in green and extra-tropics in blue. Dashed horizontal lines reflect results from time-constant models while solid skylines show time-heterogeneous rates following stratigraphic periods (shading reflect 95% HPD intervals). *Top:* fossil extinction rates for the tropics 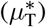 and the extra-tropics 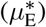. *Middle:* fossil dispersal rates from the tropics to the extra-tropics 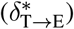 and vice-versa 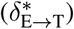. *Bottom:* fossil sampling rates for the tropics 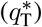 and the extra-tropics 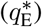. (Mastodon silhouette credits: Becky Barnes – North Dakota Geological Survey).

### Conclusions

Both our phylogenetic and paleontological results emphasize the key role that extra-tropical extinction played in shaping present-day latitudinal diversity patterns. They reject the hypotheses of an older, larger, or more speciation-prone tropical biome leading to current richness. Instead, they support the hypothesis that surviving diversity accumulated in the tropics under low extinction, dispersed out of the tropics, and suffered from frequent extinctions in the extra-tropics. These frequent extinctions did not affect species equally: some extra-tropical lineages accumulated species just as fast as those in the tropics, while others experienced negative diversification rates. These differences likely reflect species-specific traits that offer varied resilience to cope with the somewhat harsher environments of the extra-tropics, although this remains to be tested. Our results show an extraordinary correspondence in biome-dependent diversification and dispersal patterns across the four tetrapod clades, despite each clade’s idiosyncratic responses to paleoenvironmental dynamics. Markedly distinctive ecologies across clades underlie multiple routes to an overarching latitudinal variation in geographical diversification that brought about one of the most striking diversity patterns on our planet.

## Methods

### Tropical and extra-tropical definition

We use the Köppen biome classification (Köppen, 1884) to define tropicality and extra-tropicality. The Köppen biome classification is based on temperature and precipitation and their seasonal components and delimit ecologically meaningful regions (Beck et al., 2018). While there are several ways to define the tropics and extra-tropics (*e*.*g*., 23.5° south and north latitudes as somewhat arbitrary thresholds), we adopt our classification for two main reasons. Firstmost, climatic variables are the major determinants of species distributions, providing a biologically meaningful spatial delimitation when explaining species richness patterns. Climatic variables are strongly correlated with latitude, but local topographical and regional features can distort the expectation of a tropical biome, even at low latitudes (Alves et al., 2017). Secondly, our delimitation matches the spatial paleoclimatic reconstruction (see below) based also on the Köppen classification scheme to estimate tropical and extra-tropical area through deep-time.

For present-day spatial tropical delimitation, we extracted the raster data at 1 km resolution from (Beck et al., 2018), who divided the terrestrial globe into five main classes and thirty sub-types. We aggregated all tropical subtypes (*i*.*e*., “Af”:Tropical Rainforest, “Am”:Tropical Monsoon, “Aw”:Tropical Savannah) to define a tropical biome, and aggregated the remaining subtypes as extra-tropical (Beck et al., 2018). Effectively this delineates the tropics as a region where the Mean Air Temperature of the coldest month is above 18° C and that has high Mean Annual Precipitation (see (Beck et al., 2018) for details). We then reprojected the raster data into the Mollweide projection, which preserves equal-area proportions across the globe (Fig 1a).

### Terrestrial vertebrate data

We aggregated phylogenetic and spatial information for all extant species of amphibians, squamates, birds, and mammals. Phylogenetic data for amphibians was obtained from (Jetz and Pyron, 2018) (7238 total species, 4061 genetically represented), for squamates from (Tonini et al., 2016) (9755 total species, 5415 genetically represented), for birds from (Jetz et al., 2012) (9993 total species, 6670 genetically represented) and for mammals from (Upham et al., 2019) (5806 total species, 4001 genetically represented). We excluded the Tuatara species (*Sphenodon punctatus*) from the phylogeny in (Tonini et al., 2016) because Tuatara is not a squamate. We excluded marine-breeding mammal species (*i*.*e*., Cetaceans, 105 species) since terrestrial covariates are unsuited for assessing their evolutionary history. We excluded 13 species of domestic animals for they have mostly responded to human selection and have unusual human-mediated cosmopolitan biogeographic distributions, which might conflate the natural diversification dynamics, but included many human-induced recent extinct species (103 spp.). For each clade we randomly sampled 10 trees from the posterior to accommodate phylogenetic uncertainty in posterior analyses.

Spatial information was carefully vetted and collated to match the taxonomy of the phylogenetic trees, obtained from VertLife project (vertlife.org) in association with Map of Life (mol.org). Distribution data for birds followed (Jetz et al., 2012). Mammal and amphibian distribution data were based on IUCN data (for Conservation of Nature, 2016) that were modified to match taxonomically with the respective phylogenies. For squamates data was based on (Roll et al., 2017) with careful taxonomic matching with the phylogenetic tree. We performed a literature search to assign those species without spatial distribution information as either tropical, extra-tropical or widespread (species-specific references can be found in Table S1). For mammals, we note that we included recently extinct fossils, which have been sampled from few locations and might not be representative of the whole range (Table S1).

To encode species ranges as tropical, extra-tropical, or widespread, we used expert breeding ranges defined for 360 equal area grid cells (110km grain size) since they have been previously validated to have minimum (*<* 5 − 10%) false presences at such spatial grain (Hurlbert and Jetz, 2007). We then counted the number of tropical and extra-tropical 1km cells from the binary Köppen layer inside each species range to assign them as tropical, extra-tropical or widespread using 4 different ‘cut-offs’. As the most stringent cut-off, we required at least 20% of the range to pertain to either the tropical and extra-tropical area to be assigned that area; then we used 15%, 10% and 5% in decreasing order of regional exclusivity (Alves et al., 2017). Complementary analyses showed that the different cut-offs did not significantly impact our primary findings concerning diversification patterns (Fig S2) and thus we carried on using only the most stringent cut-off of 20 %, which is conservative in allocating tropical and extra-tropical taxa. Briefly, the least conservative cut-off of 5% produces a higher proportion of widespread lineages (concomitant with fewer endemics), which, in turn, increases dispersal rates from the tropics to the extra-tropics and decreases *in situ* speciation rates in the extra-tropics compared to the 20% cut-off (Fig S2).

After allocating all extant species of amphibians, squamates, birds and mammals as tropical, extra-tropical or widespread, we pruned all species from each phylogeny that were not placed using genetic data, and then estimated the sampled fraction for each of those states in the resulting trees. The resulting sampling fractions for the tropics (T), extra-tropics (E) and widespread (W) taxa are for amphibians (T = 0.520, E = 0.594, W = 0.585), for squamates (T = 0.464, E = 0.648, W = 0.569), for birds (T = 0.574, E = 0.775, W = 0.681), and for mammals (T = 0.612, E = 0.767, W = 0.686), respectively, and showcasing the overall genetic undersampling of the tropics. Our main results used only genetically sampled species to avoid biases in assuming an artificial geographic evolutionary history resulting from any unnatural placement of non-genetically sampled species (Hagen et al., 2021). Expectedly, random placement of non-genetically sampled species will spuriously yield more allopatric sister taxa than one might expect when using such large geographical regions. In any case, we compared our results using all species, obtaining similar relative differences but, as expected, with inflated rates of transitions and between-region speciation (Fig S2).

### Tip speciation rates analyses

To estimate tip speciation rates for each of the four tetrapod clades, for each of the 10 posterior tree samples, we made inference under the ClaDS model (Maliet et al., 2019) using the data augmentation implementation (Maliet and Morlon, 2022). ClaDS assumes speciation rates are constant within each branch of the tree but are inherited with a shift at speciation, thereby providing branch-specific speciation rates. We ran ClaDS using ambiguous priors on the hyperparameters under a constant turnover model for as many iterations as necessary for the Gelman-Rubin statistic (Gelman and Rubin, 1992) to become lower than 1.05 (the default stopping behaviour). We then retrieved the present-day speciation rates for each species by taking the maximum *a posteriori* estimate for each of the 10 phylogenetic trees for each clade.

To test for a latitudinal effect on speciation rates, we used Bayesian linear models corrected for phylogenetic non-independence using the MCMCglmm (Hadfield, 2010) package in R (R Core Team, 2020). We first calculated the latitudinal centroid for all species and explored the effect of absolute latitude centroid on the natural logarithm of tip-speciation rates. We then estimated if there was a difference between tropical and extra-tropical biomes in speciation rates. We ran the analyses for each clade, and for each tree for 500k MCMC iterations with a 2K burn-in using ambiguous inverse Wishart priors (expected covariance of 1 and *ν* = 2*e* − 4) and visually checked for good mixing and convergence.

### Paleoenvironment reconstructions

To quantify the tropical and extra-tropical terrestrial area through time, we used plaeoclimatic and paleoelevational reconstructions that date as far as 540 Mya with a sampling frequency of every 5 Myrs. Paleo-Digital Elevation Models, “PaleoDEMs”, were obtained at a 1° resolution from (Scotese, 2021). Similarly, for the same time stamps, we used Köppen classification reconstructions from (Scotese, 2021) with corresponding 5 Myr interval maps. These reconstructions are based solely on the five main climate classes (*i*.*e*., Tropical, Arid, Warm Temperate, Cool Temperate, and Polar), which we divided into tropical and extra-tropical. We reprojected these data into the Mollweide projection and intersected terrestrial land, by using only gridcells above sea-level, with either tropical and extra-tropical classification. Finally, we counted the number of grid cells and multiplied by their area to obtain tropical and extra-tropical area every 5 Myrs from the present back to 540 Mya (Fig 4 & Video S1). All spatial analyses where conducted with gdal (GDAL/OGR contributors, 2020) and raster package (Hijmans, 2019) for R (R Core Team, 2020). We used deep-time global Temperature estimates for the Phanerozoic eon from (Condamine et al., 2019). For both curves we used spline smoothing in R using the pspline package (Ramsey and Ripley, 2017; R Core Team, 2020). For Temperature (‘T(t)’) and biome specific Area (‘A(t)’) we also estimated their rates of change by taking the derivative from the smoothed curves (‘T’(t)’ and ‘A’(t)’, respectively). Finally, we tested diversity-dependence by using an exponential dependency with time (noted ‘ETD’) (Rabosky and Lovette, 2008).

### Effect of environment on diversification across space

We expand on current State dependent Speciation and Extinction (SSE) models to allow for environmental dependency. We call these set of models Environmental SSE models, ‘ESSE’. In general, SSE models enable inference on distinct rates for speciation, extinction and state transitions depending on a discrete state, yet for a given state, remain constant through time. (Cantalapiedra et al., 2014) enabled environmental dependency on speciation rates for MuSSE within a likelihood framework (FitzJohn, 2012), using a different mathematical formulation than the one presented here. We generalize by allowing rates of speciation, extinction and state transitions to depend on discrete states, including hidden states as in ‘HiSSE’ (Beaulieu and O’Meara, 2016), as well as time-varying covariates ***z***(*t*), whose effect on rates of speciation, extinction and state transitions is regulated by parameters *β* within a Bayesian framework.

Our ESSE model elaborates on the Geographic State Dependent Speciation and Extinction (‘GeoSSE’) model introduced by (Goldberg et al., 2011) and later enhanced by (Caetano et al., 2018) to include Hidden States (‘GeoHiSSE’) to consider more than 2 areas. In GeoHiSSE models, each area *i* and hidden state *h* has a rate of speciation (*λ*_*i,h*_) and extinction (*μ*_*i,h*_), each pair of areas *i, j* and hidden state *h* have a rate of dispersal (from *i* → *j, δ*_*i,j,h*_ *&* from *j* → *i, δ*_*j,i,h*_) and a rate of between-region (*i*.*e*., allopatric) speciation (*λ*_*i,j,h*_), and a transition rate between hidden states *h* and *l* (*ϕ*_*h,l*_; details in Supplementary Information). ESSE also extends ideas from the Feature-Informed GeoSSE (‘FIG’) model () by allowing per-area speciation, extinction or dispersal rates to vary according to time-dependent state-specific covariates, such as paleobiome size or temperature. While we enable here the most general model, in practice some parameters have to be constrained (see below), resulting in simpler models, to perform effective inference. Let *z*_*i*_(*t*) be a vector of length *p* at time *t* of some arbitrary functions that change through time (*e*.*g*., biome area through time), specific to lineages in area *i*, and let *β*_*γ,i,h*_ be a *p* × 1 matrix of the parameters regulating the effect of *z*_*i*_(*t*) on rate *γ*_*i,h*_ experienced by a lineage in area *i* with hidden state *h*. We then allow per-area speciation (*λ*) and extinction (*μ*) rates, *γ*_*i,h*_ = {*λ*_*i,h*_, *μ*_*i,h*_}, to depend on *z*_*i*_(*t*) multiplicatively, such that *γ*_*i,h*_(*t*) = *γ*_*i,h*_ exp (*z*_*i*_(*t*)^t^*β*_*γ,i,h*_), where *γ*_*i,h*_ is a baseline constant rate not influenced by timevarying covariates. For a covariate effect on dispersal rates, *δ*, the covariates affect pairwise area rates, such that *δ*_*i,j,h*_(*t*) = *δ*_*i,j,h*_ exp (*z*_*i,j*_(*t*)^t^*β*_*δ,i,j,h*_), where *δ*_*i,j,h*_ is the colonization rate from area *i* to area *j* under hidden state *h*. The likelihood calculations, the estimation of posterior marginal probabilities for ancestral states (Figs S3-6), Bayesian inference and validation details are given in the Supplementary Information (Fig S1).

### Implementation

These methods are available as part of the ‘Tapestree’ (Quintero and Landis, 2020) and ‘PANDA’ (Morlon et al., 2016) packages for Julia *>* v1.5 (Bezanson et al., 2017). The likelihood calculations were programmed using the DifferentialEquations package (Rackauckas and Nie, 2017) to perform numerical integration of the ODE functions. We use ‘meta-programming’ techniques to produce specialized code at run time given the model specified by the user to attain higher performance. Particularly, we are able to generate state specific functions, including the ODE function to be passed to the numerical solver. Our ESSE implementation allows for different combination of models that can be specified: i) designation of rate dependency (*i*.*e*., if speciation, extinction and/or dispersal rates are specified as dependent on a series of time-varying covariates); and ii) number of covariates per covariate-dependent rate (*i*.*e*., *p*); iii) number of states (*i*.*e*., |*K*|); and iv) number of hidden states (*i*.*e*., |*H*|). Furthermore, any number of logical constraints between parameters, or fixing of parameters to a determined real value can be specified. Code is available online at https://github.org/ignacioq/Tapestree.jl (*Note to reviewers: code will be merged into the master version of Tapestree and PANDA once the article is published*.).

### Empirical inference

For each tetrapod clade we performed inference on each of 10 phylogenetic trees sampled from the posterior to accommodate phylogenetic uncertainty. To gain a comprehensive picture of the diversification dynamics and aid in interpretation, we fitted progressively more complex geographic models. First we ran a geographically dependent model with time-constant rates and without hidden states (‘G’; *i*.*e*., our own implementation of ‘GeoSSE’). The second model included environmentally dependent speciation rates but still no hidden states (‘G+E’). The third model included two hidden states but no environmentally dependent rates (‘G+H’; *i*.*e*., our own implementation of ‘GeoHiSSE’). The fourth model included both environmentally dependent speciation rates and two hidden states (‘G+E+H’). For all models all parameters varied freely (including transition rates among hidden states), except for constraining the local extinction rate to be equal to the global extinction rate (see Supplementary Information). We used uninformative exponential priors, Exp(0.1), for all rate parameters and uninformative Gaussian priors for *β*, N(0, 10). Each run consisted on 3 Metropolis Coupled Markov Chain Monte Carlo chains and Δ*T >* 0.5 to guarantee as much as possible good convergence behaviour over the complex posterior surface. This resulted in 6 diversification scenarios (5 environmentally dependent and one under rate constancy), including or not hidden states, yielding 12 models, each ran on 10 posterior trees for each of four tetrapod clades, that is, a total of 480 model fits (aside from simulations and runs to appraise sources of bias; Figs S8 & 9).

### Bayes Factors and model comparison

We used Bayes Factors (BF) to test the relative support of processes acting differently across regions in generating the observed richness patterns. Given that our set of models are nested, we used the Savage-Dickey ratio to estimate BF (Verdinelli and Wasserman, 1995; Suchard et al., 2001). As this method is reliable only when evaluating at most two-dimensions of the posterior distribution estimated from MCMC samples (because of the curse of dimensionality), we use it only to compare models without hidden states. When the priors for the parameter of interest are independent (as is our case), the BF is given by:

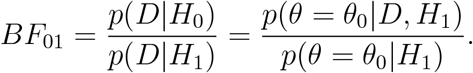

That is, the BF in support of model *H*_0_ in comparison to *H*_1_ is proportional to the posterior probability density divided by the prior probability at *θ*_0_ for the *H*_1_ model. First, we test the relative Bayesian support between models with either equal speciation or equal extinction between tropics and extra-tropics against a model with different rates per region. To this end, we test whether the derived parameter *dλ* = *λ*_*T*_ − *λ*_*E*_ is significantly different from 0. The prior for the speciation rates of each region are independent exponential distributions *λ*_*i*_ ∼ Exp(*r*_*i*_), which induce the prior probability 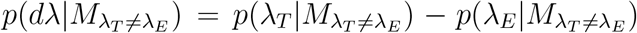 such that (for derivation see Supplementary Information):

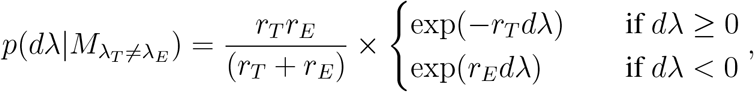

which, for the prior used for rates in our analysis of *λ*_*i*_ ∼ Exp(0.1), yields 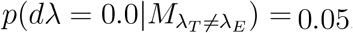. To estimate 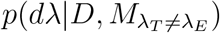, we simply use the output of the MCMC to empirically estimate the posterior distribution of *dλ* and use the polspline package (Kooperberg, 2020) in R (R Core Team, 2020) to approximate the density function and evaluate it at *dλ* = 0. The same procedure was employed to test for equal extinction rates. Similarly, to compare models with and without time-varying covariates, we evaluated the prior at *β*_*T*_ = 0 and *β*_*E*_ = 0 and the joint posterior distribution of (*β*_*T*_, *β*_*E*_) at (0,0) to estimate the Bayes factors. To estimate the joint posterior we used the np package (Hayfield and Racine, 2008) for R (R Core Team, 2020), which applies the kernel unconditional density method from (Li and Racine, 2003). With these ratios, we can then straightforwardly compare BFs among the different covariates to select the best models. Because we used the same priors on all models, the BFs estimated through the Savage-Dickey

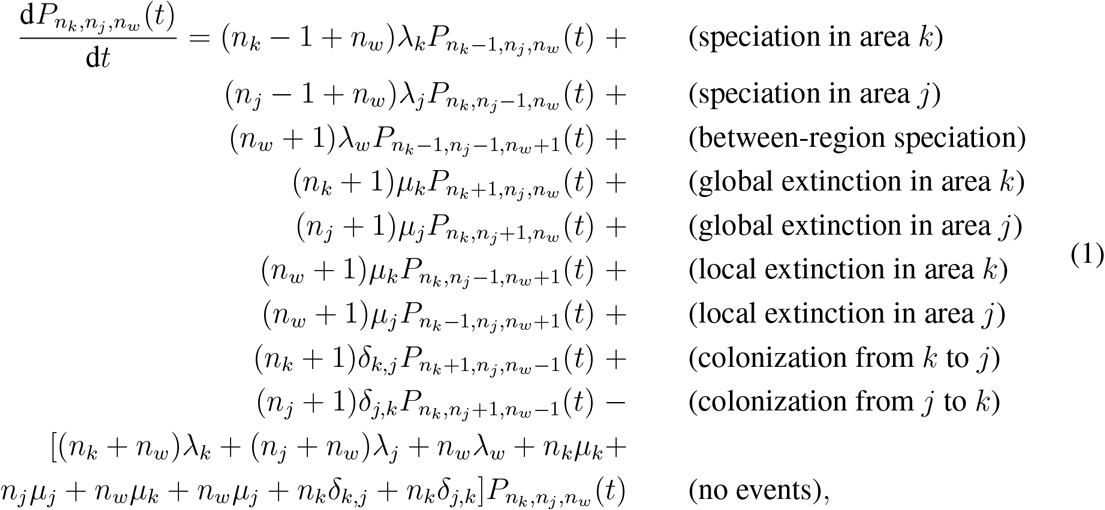

ratio yield the same result as using the Harmonic Mean (HM) approximation for the marginal likelihood constant. While the HM method usually overestimates the marginal likelihood and can misbehave when the posterior is not readily integrable (Xie et al., 2011), in our particular case, given that the models are either nested or have the same number of parameters, leading to similar overestimations, it should be a reasonable approximation for model comparison. We then used HM to estimate Bayes Factors for moddels with hidden states, for which we cannot reliably estimate their Savage-Dickey ratio. Nonetheless, we manually checked for explanatory power from the different covariates by analyzing how the posterior distributions of *β* differ from 0. Finally, we further tested model fit by performing simulations from the best performing models, as follows.

### Predictive power

We test if our best-fitting models also have good power to predict current latitudinal patterns of species richness across tetrapods. We aim to compare the observed numbers of tropical, extra-tropical and widespread species to regional richness probability distributions (RPDs) under our four models: G, G+E, G+H and G+E+H. Because no analytical solution is available for RPDs under these models, and approximating them entirely with simulations resulted in a non-negligible fraction of cases exceeding the guard of a maximum of 100,000 total simulated lineages put in the first place to avoid high computational costs, we used a combination of simulations and parametric approximation of the RPD.

We performed 500 simulations for each of the 10 phylogenetic trees for each clade for the best four models: G, G+E, G+H and G+E+H. To this end, we sampled randomly from the posterior parameter samples and corresponding ancestral estimate to start the simulation. Because we condition on both crown lineages surviving to the present, we started the simulations with a speciation event and ensured that none of the crown lineages went extinct. For each of the 10 trees for each of the clades for each model, we ran the simulations for the full tree height, stopped the simulations that hit the guard, and recorded information on the number of tropical, extra-tropical and widespread species generated for both the simulations that hit the guard, *π*_*u*_, as well as the ones that finished before, *π*_*d*_. Note that the guard considers only the total number of alive lineages without regard to the current distribution of lineages across states, so even for simulations that hit the guard, the number is not the same across states when stopping the simulation.

Next, we assumed that the Negative Binomial (NB, a generalization of the Geometrical distribution) provides an adequate representation of the RPDs. This was justified by the analytical formulas for the RPD under a constant birth-death model with survival conditioning on both crown lineages which induce a zero-inflated geometric distribution (Kendall, 1948; Stadler, 2012), because the sum of two NB distributions also follow a NB distribution, and given our own validation following the numerical solutions of the RPD under the G model (see below). We can then use the information on the *π*_*d*_ regional species richness and the *π*_*u*_ simulations that exceed the guard to obtain the maximum likelihood parameters of the NB distribution. This results in maximizing the following likelihood function

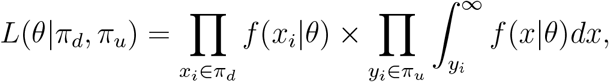

where *f* represents the probability distribution for the Negative Binomial distribution and *θ* represents its parameters (mean and size). We use the Non-Linear Minimization function ‘nlm’ in R (R Core Team, 2020) to arrive at the maximum likelihood estimates.

We developed numerical solutions for the RPDs in a two area geographic SSE process (our ‘G’ model) by first deriving their coupled dynamical equations. Let *λ*_*k*_ and *λ*_*j*_ be the *in situ* speciation rate for area *k* and *j, λ*_*w*_ be the between-region speciation rate, *μ*_*k*_ and *μ*_*j*_ be the extinction rate for area *k* and *j*, and *δ*_*k,j*_ and *δ*_*j,k*_ be the colonization rate for a lineage in area *k* to colonize area *j* and vice versa. To estimate the distribution of the number of species after time *t* under this model, we expand on the coupled dynamical equations for the joint probability of having *n*_*k*_ species in state *k* (*i*.*e*., endemic to area *k*), *n*_*j*_ species in state *j* (*i*.*e*., endemic to area *j*) and *n*_*w*_ species in state *w* (*i*.*e*., widespread), after time *t* in Eq. 1, for *n* = (*n*_*k*_ + *n*_*j*_ + *n*_*w*_) ≥ 1.

Because we only consider processes that do not go extinct, the probability for *P*_0,0,0_(*t*) = 0 throughout. Finally, to arrive at crown conditioning, that is, start the process with two lineages that do not go extinct following a speciation event, we simply make the appropriate sum across lineages for two independent processes starting with one lineage with their corresponding state. We numerically solve these three-dimensional joint RPD by assuming a very small time step and integrating after some time *t*. Estimating this joint RPD requires the integration over three dimensional matrices for every small step, making computation unfeasible for more than 200 species. Overall, we find that the NB parametric approximation accurately models the RPD, particularly as the expected number of species increases (Fig S10). Moreover, it generally yields marginally tighter (more conservative) quantiles than the RPD, resulting in robust RPD quantile estimates. We use the NB to estimate RPD quantiles per tree, per clade, per region and compare it to the empirical estimates. These numerical solutions are included in the software package.

### Biogeographic fossil analyses of Mammals

To test for congruent historical evolutionary dynamics from both neontological and paleontological evidence, a clade with both well-known fossil record and complete phylogenetic tree is required. We thus focused on Mammals, which, during the Cenozoic, remain the best sampled major tetrapod radiation in the fossil record (Mannion et al., 2014; Davies et al., 2017). We downloaded Mammal fossil occurrence data from the Paleobiology Database (PBDB) on July 2022 (Database, 2022) for all mammals (*i*.*e*.,’Class’ = ‘Mammalia’) excluding Cetaceans, since we work with terrestrial regions. We first vetted the resulting database by removing occurrences with a reported interval larger than 5 My for occurrences younger than 23.04 Mya, larger than 10 My for occurrences younger than 66.02 Mya, and larger than 15 My for all other occurrences. We then discarded all occurrences without a defined genus classification. Following common practice in paleontology, we performed all subsequent analyses at the genus level, which are less contingent on preservation and sampling biases, correlate well with species-level patterns of diversity, and better reflect underlying diversification patterns (John Sepkoski, 1998; Jablonski and Finarelli, 2009). To incorporate dating uncertainties, we produced 10 randomized datasets by sampling uniformly within the temporal range of each fossil occurrence (Silvestro et al., 2016). We then used the tropical and extra-tropical paleoenvironmental reconstructions to assign each fossil occurrence as tropical or extra-tropical according to their timing and paleocoordinates, and allowing a buffer diameter of 100 km, following the grain from the spatial reconstructions. This resulted in 74, 722 fossil occurrences allocated across the two regions, including occurrences older than the Cenozoic (Table S4). While we only considered the Cenozoic, including these occurrences is important for better estimation of sampling rates across genera that survived the K-Pg mass extinction.

We then used the Dispersal-Extinction-Sampling (‘DES’) model (Silvestro et al., 2016), which uses MCMC to estimate per-area dispersal, extinction, and sampling rates, while accounting for several sources of uncertainty such as sampling and preservation, and outperforms other methodologies that infer past diversification dynamics. We ran two separate models on the same dataset: first we constrained dispersal and extinction rates to be constant across the Cenozoic for each area, and then we allowed for time-heterogeneity by allowing rates to change according to stratigraphic ranges. For both models we allowed time-heterogeneous sampling rates. We ran each of the 10 replicates for 2 × 10^5^ iterations, and monitored convergence to then remove 10^5^ iterations as burn-in. For the final results we combined the 10 replicates, which showed very little variation between them.

## Acknowledgments

We thank Christopher R. Scotese for sharing the palaeogeographic information. We thank Daniele Silvestro, Juan L. Cantalapiedra and Juan D. Carrillo for help with fossil analyses. We thank Jérémy Andréoletti and the Morlon lab in general for feedback on an earlier draft. We are grateful to the Map of Life team, in particular Yanina Sica and Ajay Ranipeta, for support in preparing the spatial biodiversity data. This project has received funding from the European Union’s Horizon 2020 research and innovation programme under the ERC CoG PANDA to HM and Marie Skłodowska-Curie grant agreement No 897225 for IQ. MJL was supported by NSF-DEB grant 2040347. WJ acknowledges support from NSF VertLife grant DEB 1441737.

